# Two-stage adaptation of inhibition mediates predictive sensitization in the retina

**DOI:** 10.1101/214130

**Authors:** David B. Kastner, Yusuf Ozuysal, Georgia Panagiotakos, Stephen A. Baccus

**Author notes:** Present Address: Department of Psychiatry, University of California, San Francisco, 401 Parnassus Avenue, San Francisco, CA, 94143 USA. Present Address: Google Inc., Mountain View, CA 94043, USA. Present Address: Broad Center of Regeneration Medicine and Stem Cell Research, University of California, San Francisco, 35 Medical Center Way, San Francisco, CA 94143, USA. Lead contact: Stephen A. Baccus.

## Abstract

A critical function of the nervous system is the prediction of future sensory input. One such predictive computation is retinal sensitization, a form of short-term plasticity seen in multiple species that elevates local sensitivity following strong local stimulation. Here we perform a causal circuit analysis of retinal sensitization using simultaneous intracellular and multielectrode recording in the salamander. We show, using direct current injection into inhibitory sustained amacrine cells that a decrease in amacrine transmission is necessary, sufficient and occurs at the right time and manner to cause sensitization in ganglion cells. Because of neural dynamics and nonlinear pathways, a computational model is essential to explain how a change in steady inhibitory transmission causes sensitization. Whereas adaptation of excitation removes an expected result in order to transmit novelty, adaptation of inhibition provides a general mechanism to enhance the sensitivity to the sensory feature conveyed by an inhibitory pathway, creating a prediction of future input.

## 1 Introduction

Understanding how individual components of a neural circuit contribute causally to a biological function presents a challenging problem. Within a neural circuit, signals travel through serial connections and parallel pathways through a diversity of cell types. The components of those circuits often have nonlinear and interdependent effects, meaning that the effects of individual mechanisms must be considered in the context of a particular computation. Consequently, the mechanisms of even well studied neural computations, such as the receptive fields of orientation selective neurons in the primary visual cortex (Hubel and Wiesel, 1962; Martinez, 2011), remain incompletely understood.

This problem is analogous to the classical problem of demonstrating that a molecular mechanism causes a developmental process or that a pathogen is responsible for a disease. The typical goal is to establish simple causation by blocking or activating a particular molecular component and measuring the effect on a downstream process.

The problem of distinguishing a mediator from a modulator has challenged studies of molecular processes for long term neural plasticity (Sanes and Lichtman, 1999; Lisman et al., 2003). In neural circuits, this problem is compounded because of the pervasiveness of parallel processing and nonlinear dynamics. Furthermore, this problem is underappreciated with regards to neural circuits, where it has become common to draw conclusions based on perturbations alone such as using optogenetics, yielding mistaken interpretations (Otchy et al., 2015; Phillips and Hasenstaub, 2016). Consequently, no currently accepted approach of experimentation and computational analysis exists to characterize the causal role of a particular cellular or molecular mechanism to a dynamic neural computation, especially in the face of complex and unknown nonlinear effects.

Circuit computations—those that arise not by the action of a single cell but by the interaction of multiple neurons in a circuit—present a particularly difficult challenge for mechanistic inquiry because of the need to study the intact circuit. Yet circuit computations also provide an opportunity for understanding because of the ability to perturb neurons in the circuit as it operates. One such circuit computation is retinal sensitization, a process seen in multiple species that elevates local sensitivity following strong local stimulation (Kastner and Baccus, 2011; Nikolaev et al., 2013; Kastner and Baccus, 2013). Theoretical analyses and experiments have indicated that this elevation of sensitivity embodies a prediction that a target stimulus feature will be present in the future in that same region (Kastner and Baccus, 2013). The prediction of future sensory input is an important overall function of the nervous system, (Montague and Sejnowski, 1994; Schultz et al., 1997), yet the mechanisms of such computations are generally unknown. Although transmission from GABAergic amacrine cells is required for sensitization (Kastner and Baccus, 2013), the slow action of pharmacological manipulations can cause compensatory actions in the circuit, making a conclusion uncertain (Cook et al., 2000).

Here we use simultaneous intracellular and multielectrode recording in the salamander retina to establish quantitatively the causal role of a class of inhibitory interneurons in the computation of sensitization. We take the generalizable approach of first measuring the responses and transmission of an interneuron to create a computational model of the input and output pathways of the interneuron and of the parallel pathways that do not travel through the cell. We find that a decrease in transmission in two stages of an inhibitory pathway causes sensitization. Both the input to and the output from sustained amacrine cells decreases during a localized sensitizing stimulus. We further use current injection to replay the input experienced by an individual amacrine cell, and found this input was sufficient to cause sensitization. In addition, the measured changes in amacrine cell transmission were sufficient to cause ganglion cell sensitization. Finally, a computational model characterizes the change in steady inhibitory transmission, and shows that it accounts quantitatively for the observed sensitivity changes.

Behavioral sensitization has been attributed to changes in excitatory synaptic strength or intrinsic excitability (Dale et al., 1988). At the circuit level, adaptation of excitation underlies the subtraction of a prediction, by diminishing the expected result of the circuit and thereby transmitting a signal enriched for novelty (Berry et al., 1999; Hosoya et al., 2005; Schwartz et al., 2007b,a; Palmer et al., 2015). We find that adaptation of an inhibitory pathway provides a general mechanism for the presence of a sensory feature to subsequently cause an enhanced sensitivity for that feature, creating a prediction of future input.

## 2 Results

In the case of the short term plasticity process of retinal sensitization, previous results have shown that GABAergic transmission is necessary for sensitization in the salamander retina. However, these results cannot distinguish between the requirement for only a constant level of inhibition and a dynamic change in inhibition as would occur through changes in inhibitory transmission. Furthermore, such results neither show that any change in inhibition occurs during sensitization, nor that a change in inhibition is sufficient for sensitization.

### 2.1 A sensitizing stimulus creates an afterhypolarization in amacrine cells

As signals are processed by the retina from photoreceptors to ganglion cells, the effect of a particular interneuron such as an amacrine cell can be logically divided into two stages: 1) the input to the amacrine cell, consisting of the circuitry that leads from photoreceptors to the amacrine cell; and 2) The output from the amacrine cell, comprising the circuitry that leads from the amacrine cell to the ganglion cell. Understanding the contribution of an interneuron to the processing of the entire circuit requires measurement of both the membrane potential response of the neuron—reflecting its input—and the effects of its output on the circuit (Lehky and Sejnowski, 1988).

We first sought to determine whether this first stage—the amacrine cell input stage—exhibited any change in processing that was correlated with the dynamics of sensitization. To do this, we simultaneously recorded intracellularly from a sustained Off-type amacrine cell and extracellularly from multiple nearby ganglion cells, all while presenting to the retina a visual stimulus that causes sensitization by increasing and then decreasing the temporal contrast (Figure 1 – D). We focused on fast Off type adapting and sensitizing ganglion cells, two salamander cell types that form independent mosaics across the retina (Kastner and Baccus, 2011). Both cell types sensitize, although adapting cells have stronger adaptation in the receptive field center that cancels some sensitization (Kastner and Baccus, 2013).

**Figure 1:**
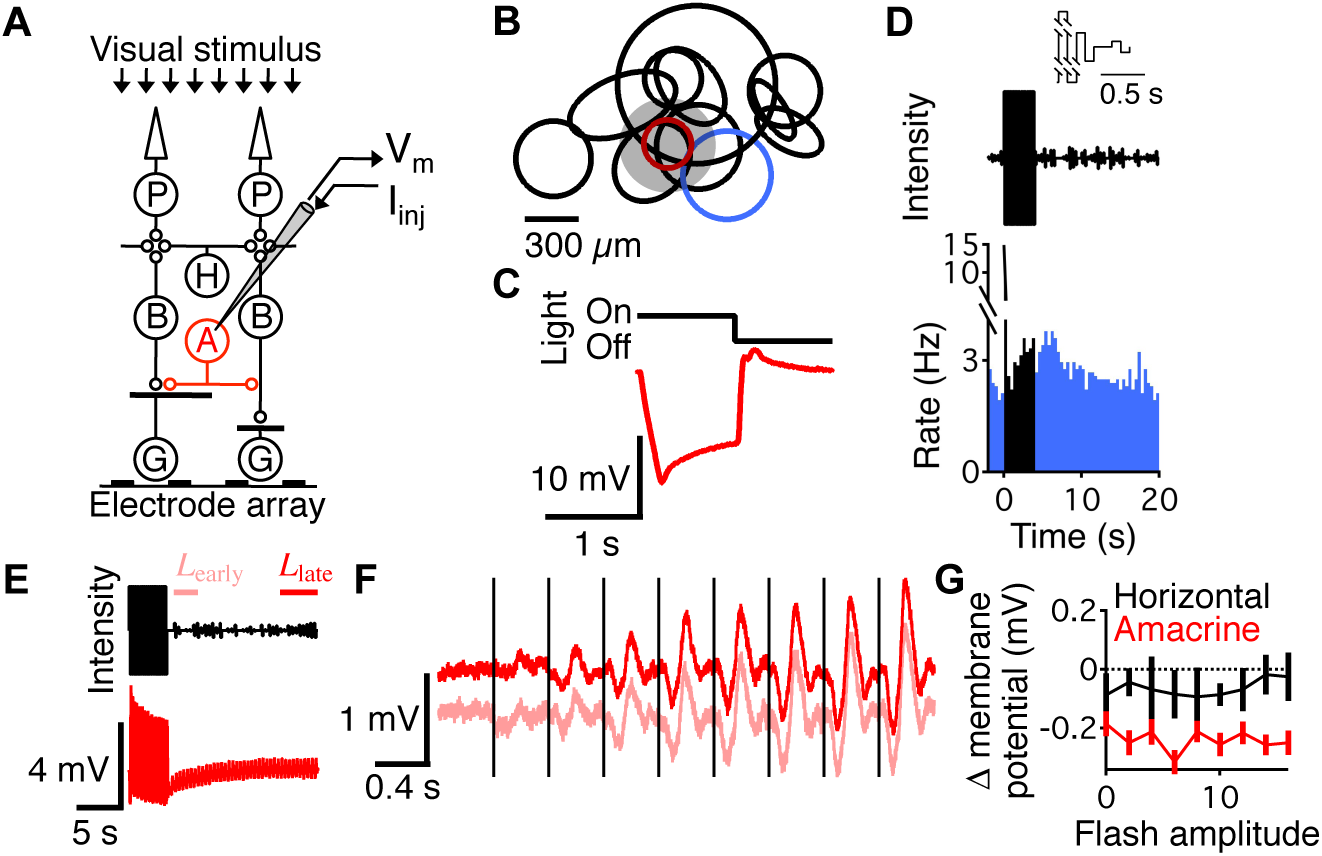
Amacrine cells hyperpolarize after a sensitizing stimulus. (A) Experimental setup for simultaneous intracellular and multielectrode recording. (B) Spatial receptive fields of an amacrine cell (red) and ganglion cells recorded simultaneously. Gray shaded region shows the localized spot that flickered at high contrast to generate sensitization. (C) Average response of a sustained Off amacrine cell to a 0.5 Hz flashing stimulus. (D) Visual stimulus (top) and the response (bottom) of an example ganglion cell (blue receptive field from panel B) showing sensitization as an elevated response to low contrast (colored trace) following high contrast. High contrast was 100% Michelson contrast, and low contrast was composed of 9 different 200 ms biphasic flashes (inset) randomly interleaved. The high contrast was only presented over the receptive field of the amacrine cell, and the rest of the image was uniform field low contrast. (E) Visual stimulus (top) and the response of an example amacrine cell (bottom, red receptive field from B), which became rectified at high contrast as indicated by larger depolarizing fluctuations. (F) Average membrane potential of the amacrine cell shown in E to the nine different low contrast flashes. Time intervals for *L*_*early*_ and *L*_*late*_ are shown above the stimulus in E, and although the different flashes were randomly interleaved, they are shown here concatenated in order of increasing strength, with vertical lines indicating the start of each flash. (G) Average difference across the duration of the flash in membrane potential between *L*_*late*_ and *L*_*early*_ for amacrine (n = 33, red) and horizontal (n = 4, black) cells. Negative values indicate that the cell was more hyperpolarized during *L*_*early*_ compared to *L*_*late*_. Error bars indicate s.e.m.

We tested whether individual sustained Off amacrine cells (Figure 1), which in the salamander are GABAergic (Yang et al., 1991), changed their response properties to a localized flashing spot stimulus that sensitizes ganglion cells. To localize the effect to the single amacrine cell we were manipulating, we created an online map of the receptive field of the amacrine cell (Figure 1) using a white-noise checkerboard stimulus. This allowed us to place a flickering high contrast spot over the amacrine cell and then measure its membrane potential when the spot changed from high to low contrast (Figure 1). Such a localized high contrast also sensitizes more ganglion cells than does a uniform field stimulus because of the center-surround spatial antagonism of adaptation and sensitization (Kastner and Baccus, 2013).

Sustained Off amacrine cells showed a dynamic response that correlated with sensitization—they adapted to the contrast of the focal stimulus by hyperpolarizing during sensitization (Figure 1 – G). Their membrane potential hyperpolarized immediately after the transition from a high to a low contrast, *L*_*early*_, compared to later during low contrast, *L*_*late*_. This hyperpolarization was similar to an afterhyperpolarization seen in ganglion cells and cortical neurons undergoing contrast adaptation (Carandini and Ferster, 1997; Manookin and Demb, 2006). Although sustained Off amacrine cells do not adapt to uniform-field Gaussian stimuli (Baccus and Meister, 2002), this stronger, more focal stimulus did generate adaptation. Additionally, we observed that the membrane potential response to the focal stimulus was more strongly rectified than for previous measurements to a full field stimulus, meaning that depolarizing fluctuations were larger than hyperpolarizations (Figure 1). This rectified response causes a change in the mean membrane potential at high contrast. A previously described accurate model of contrast adaptation indicates that a change in the mean signal level will cause adaptation, likely by causing synaptic depression in the bipolar cell inputs to the cell (Ozuysal and Baccus, 2012). In contrast, horizontal cell responses were not rectified and did not hyperpolarize following high contrast (Figure 1), indicating that they do not adapt to contrast, consistent with previous observations (Baccus and Meister, 2002).

### 2.2 Amacrine transmission changes during sensitization

We then measured whether during sensitization there were changes in processing in the transmission stage from the amacrine cell—the amacrine cell output stage—which is the stage between the interneuron and ganglion cells. To test whether amacrine transmission changed during sensitization, we first measured the firing rate of the ganglion cell as a function of the amplitude of a flash of light—the ganglion cell light response function, *G*(*s*). Then we measured how *G*(*s*) changed when we delivered either depolarizing or hyperpolarizing pulses (±500*pA*) to the amacrine cell (Figure 2 – B, see Experimental Procedures). The ganglion cell response curve, *G*(*s*), was approximately sigmoidal and changed when current pulses were injected into the amacrine cell.

**Figure 2:**
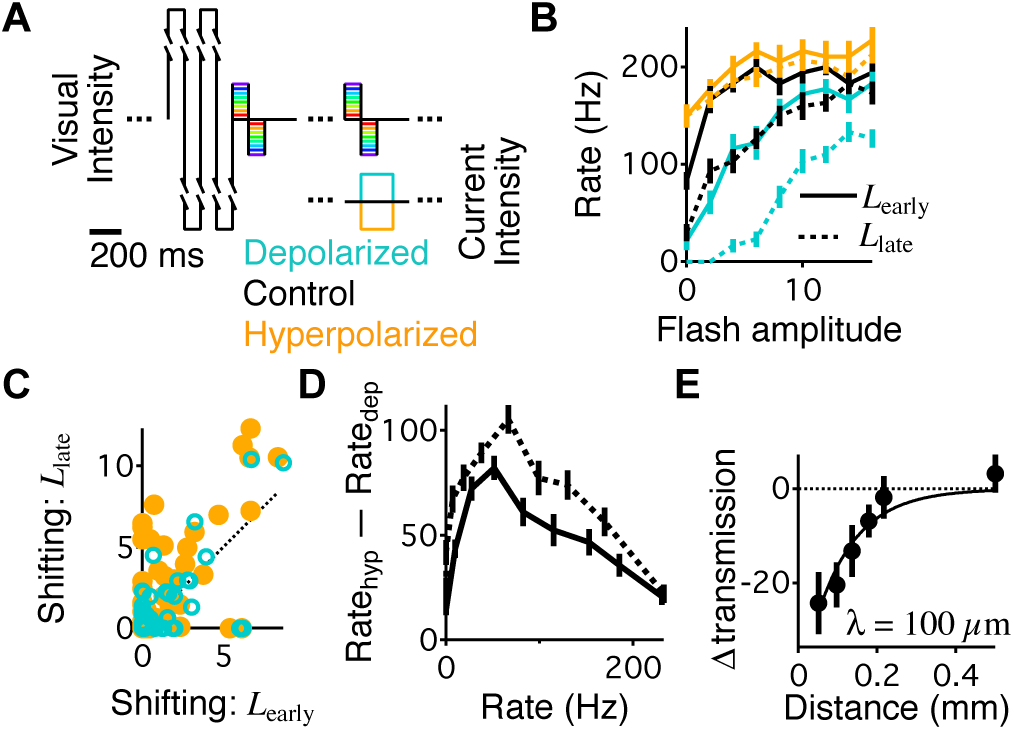
Amacrine transmission decreases during sensitization. (A) Visual stimulus alternating between high and low contrast. High contrast was 100% Michelson contrast, presented only over the amacrine cell receptive field center, and low contrast, presented everywhere, was composed of 9 different flashes randomly interleaved. Current pulses (0.5 nA) were precisely delivered for 200 ms starting when the visual stimulus flash went from on to off. Current was only delivered during the flashes occurring during *L*_*early*_ (0.8 – 3.2 s) and *L*_*late*_ (16 – 20 s) after the high contrast, and all three conditions (hyperpolarizing current, depolarizing current, and control) were randomly interleaved at each stimulus presentation. (B) Average response of an example ganglion cell to the 9 different visual flashes during the 6 conditions of the experiment. (C) The shift of the ganglion cell response curve in response to amacrine cell hyperpolarization (closed circles, orange) and depolarization (open circles, teal) during *L*_*early*_ (abscissa) and *L*_*late*_ (ordinate) (n = 51 cell pairs, 23 amacrine cells). These values were computed as the horizontal shifting parameter *µ* that captured the effect of current injection on the ganglion cell light response curve. (D) Average difference between the two current conditions in *G*(*s*) as a function of the response of the ganglion cell, ∆*G*_*A*_(*G*). Results are from all amacrine-ganglion cell pairs within 200 *µ*m of each other (n = 60 cell pairs, 24 amacrine cells). (E) Change in transmission as a function of distance between the amacrine and ganglion cell, averaged across all pairs (n = 95 cell pairs, 26 amacrine cells). The change in transmission is defined as the average difference between the effect of current on the ganglion cell firing rate as a function of the particular ganglion cell firing rate during control conditions (∆*G*_*A*_(*G*)_*early*_ − ∆*G*_*A*_(*G*)_*late*_) for each cell pair. A negative change in transmission indicates diminished transmission during *L*_*early*_ compared to *L*_*late*_.

Because amacrine cell current injection altered the ganglion cell light response curve, we sought to measure whether this effect changed during sensitization. Light responses were analyzed from ganglion cells within 200 *µ*m of the amacrine cell. To measure how the effects of amacrine transmission changed during sensitization, we measured the shift of *G*(*s*) caused by amacrine cell hyperpolarization during *L*_*early*_ and *L*_*late*_ (Figure 2). The effect of amacrine hyperpolarization was 1.74 ± 0.32 during *L*_*early*_, and 2.92 ± 0.47 during *L*_*late*_ indicating that amacrine transmission was 40% smaller during sensitization (*p <* 0.04). The effect of amacrine depolarization (1.25 *±* 0.28 during *L*_*early*_and 1.36 ± 0.32 during *L*_*late*_) was less than half of that of hyperpolarization, consistent with previous observations that sustained amacrine cells release a strong tonic inhibition and mainly act by hyperpolarization and disinhibition of ganglion cells (Manu and Baccus, 2011). Consequently, the magnitude of the effects from depolarization (Figure 2) was not significantly different across the population between *L*_*early*_ and *L*_*late*_, reducing by 9%. Thus it appeared from this simple analysis that the effect of amacrine transmission on the ganglion cell light response was smaller during *L*_*early*_ than *L*_*late*_.

A critical issue in identifying which neurons contribute to a computation in a parallel and nonlinear circuit is whether the effect of one pathway is incorrectly assigned to another. We therefore performed a further analysis to verify that the changing effect of amacrine current injection arose due to plasticity of the transmission of that cell, as opposed to being an apparent effect from unrelated plasticity elsewhere in the circuit. Even with a simple sigmoidal response curve, the apparent effect of a neuron *a* when probed by current injection can change even if the transmission of that neuron remains constant (Figure S3). This effect can occur due to a shifting of the response curve caused by changes in an independent pathway *b*. Because the effects of *a* are delivered to a different part of the nonlinear response curve, the measured effects of *a* can change even when all changes in the circuit are due to neuron *b*.

To rule out this source of an apparent change in transmission, we examined ∆*G*_*A*_ not as a function of different light levels, but as a function of different ganglion cell firing rates, *G*. We found that during *L*_*early*_, a smaller change in firing rate occurred overall (Figure 2). This decreased effect of current injection was also apparent when we controlled for the shifting ganglion cell response curve by comparing ∆*G*_*A*_ at the same ganglion cell firing rates during *L*_*early*_ and *L*_*late*_ (Figure 2 & E). Thus, the decrease in change in firing rate was not because the effect was applied at different points of the ganglion cell firing rate curve.

The decrease in amacrine transmission during *L*_*early*_ decayed as a function of distance between amacrine and ganglion cells with a space constant of *λ* = 100 ± 20*µ*m (Figure 2E). This increase in transmission from *L*_*early*_ to *L*_*late*_ is consistent with a recovery from synaptic depression, a process that exists in amacrine cells (Sagdullaev et al., 2011). These results show that given the nonlinear properties of the circuit, amacrine responses and transmission are correlated in time with sensitization.

### 2.3 Adapting inhibitory transmission in a model of sensitization

In order to develop a test for whether changes in amacrine transmission mediate sensitization, we extended a previous model that uses an adaptive component that represents the properties of synaptic vesicle release (Ozuysal and Baccus, 2012). We recorded intracellularly from a sensitizing ganglion cell responding to a uniform field stimulus. Under these conditions, sustained amacrine cells do not adapt in terms of their light response (Baccus and Meister, 2002) as opposed to what occurs in response to a spot stimulus (Figure 1E – G), allowing us to focus on how adaptation in sustained Off amacrine cell transmission (Figure 2) causes sensitization.

The pathways of the model consisted of a linear temporal filter, a static nonlinearity and a first order kinetic model of the type used to capture chemical reactions, such as synaptic vesicle recycling. This Linear-Nonlinear-Kinetic (LNK) model accurately captures the membrane potential fluctuations of adapting neurons, as well as all of their adaptive properties (Ozuysal and Baccus, 2012). The inhibitory pathway inhibited the excitatory pathway prior to its nonlinearity (Figure 3B), consistent with presynaptic inhibition onto bipolar cell terminals as has been observed in the synaptic terminals of zebrafish during sensitization (Nikolaev et al., 2013), with the threshold in the excitatory pathway corresponding to the threshold of the voltage-dependent calcium channel (Mennerick and Matthews, 1996).

**Figure 3:**
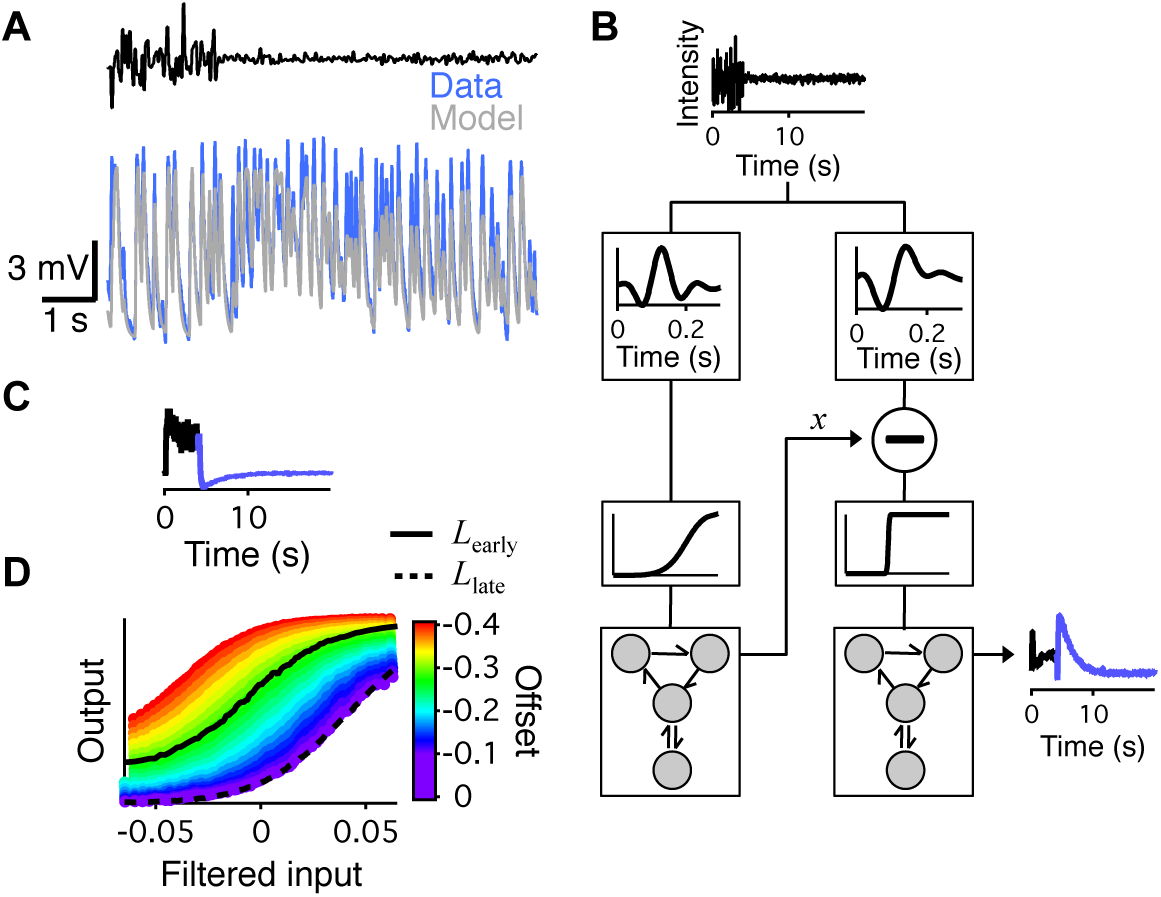
Computational model of sensitization from reduced inhibition. (A) Intracellular recording from a sensitizing ganglion cell responding to uniform field Gaussian white noise at the transition between 35% and 5% contrast, compared to the fit of the model in panel B. The membrane depolarization that underlies sensitization is evident after the transition. (B) Two pathway Linear-Nonlinear-Kinetic (LNK) model (Ozuysal and Baccus, 2012), each pathway consisting of a linear temporal filter, a static nonlinearity and a first order kinetic block (see Experimental Procedures). The inhibitory pathway (left) delivers its output prior to the threshold of the excitatory pathway (right) representing presynaptic inhibition (Nikolaev et al., 2013; Kastner and Baccus, 2013). (C) The output of the inhibitory pathway at the point labeled *x* in B. Blue indicates low contrast. (D) The nonlinearity of an LN model (different from the linear and nonlinear component of the LNK model) at different values of offset for the inhibitory pathway. Solid and dashed lines show the measured nonlinearities of an LN model for *L*_*early*_ and *L*_*late*_, respectively.

This model captured sensitization, including the millisecond timescale membrane potential fluctuations and the depolarization and shift in threshold during *L*_*early*_ (Figure 3A and 3B). We examined the output of the inhibitory pathway, and found that the mean decreased during *L*_*early*_ (Figure 3C). One of the fundamental insights derived from the use of the LNK model for contrast adaptation was that no part of the circuitry directly measures the contrast of the input. Rather, the synapse—represented by the kinetics block—always adapts to the mean, but because the signal is rectified, the mean of the signal reflects the stimulus contrast (Ozuysal and Baccus, 2012). Because of the special role played by the mean of the input to the synapse, we tested whether the mean value of the transmission from the inhibitory pathway could produce all of the effects of sensitization. When we manipulated the mean of the inhibitory pathway in the model, we changed the overall response curve of the cell smoothly between the values observed during *L*_*early*_ and *L*_*late*_, changing the threshold, offset and slope of the output response function. The model suggests that sensitization could be caused by a decrease in inhibitory transmission, which then influences the overall sensitivity of the circuit.

### 2.4 Relative contribution of adaptation from amacrine input and transmission

To estimate the relative contribution to sensitization of the two stages of decreased inhibition—decreased amacrine response (Figure 1) and transmission (Figure 2)—we presented the model of Figure 3 with the stimulus used to record the changes in the amacrine cell. Although the model only included adaptation in amacrine transmission, we added in the measured offset the amacrine membrane potential (Figure 1 E – G). We found that the measured membrane potential change from adaptation in sustained Off amacrine cells (Figure S4A), could only account for a fraction of the full magnitude of sensitization (Figure 4). Our analysis indicated that the adaption at the level of the amacrine cell membrane potential contributes less than half of the change in inhibition necessary for sensitization and that the adaptation at the level of the amacrine cell transmission contributes greater than half of the change in inhibition necessary for sensitization.

**Figure 4:**
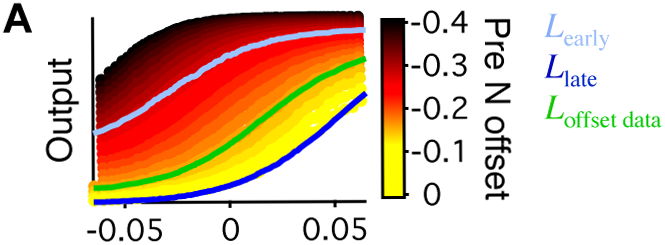
Relative contribution of amacrine cell membrane potential plasticity for sensitization. (A) Nonlinearities of LN model fits to the output of the LNK model from Figure 3 and the stimulus composed of a Gaussian white noise 3.9% low contrast preceded by a 50% high contrast (see Experimental Procedures). The light blue curve is the response of the model during *L*_*early*_. The dark blue curve is the response of the model during *L*_*late*_. The green curve is the response of the model to low contrast alone when applying an offset to the model in the inhibitory pathway following the linear filter. The offset applied matched the average change in membrane potential measured in amacrine cells during sensitization (Figure 1G, see Experimental Procedures). The range of colored curves in the background are responses of the model to low contrast alone with a range of offsets applied to the inhibitory pathway following the linear filter.

### 2.5 Changes in amacrine output can reproduce or cancel sensitization

Previous work has shown pharmacologically that inhibition is necessary for sensitization (Kastner and Baccus, 2013; Nikolaev et al., 2013). This finding, however, does not distinguish between a permissive role of inhibition, such as setting the baseline threshold of ganglion cells, and our hypothesis that changes in amacrine inhibition mediate sensitization. To test the causal effect of changes in amacrine transmission on sensitization, we analyzed whether the effect from current injection into amacrine cells was one that could cause the specific changes in the visual response of sensitized ganglion cells, and not simply a change in average firing rate. For these analyses we used the same data from Figures 1 & 2 where we presented a flashing visual stimulus to the retina that allowed for the rapid measurement of the ganglion cell response as a function of light intensity, *G*(*s*).

A change from one sigmoid to another can be accounted for by four parameters that capture: 1) a horizontal shifting of the curve along the light intensity axis, 2) a horizontal scaling, which is a change in the slope of the curve without changing the maximum, 3) a vertical shift in the curve, and 4) a vertical scaling that changes the maximum response. However, we found that only two parameters were needed to capture 79.9 ± 3.2% (hyperpolarizing) and 82.3 ± 3.2% (depolarizing) of the effects of amacrine current injection, *µ*, a horizontal shifting of *G*(*s*) along the light intensity axis, and *ν*, a horizontal scaling along the light intensity axis (Figure S2A & S2B, see Experimental Procedures).

To determine whether a decrease in amacrine cell transmission was sufficient to create sensitization, we assessed, for each cell pair, whether the effects of hyperpolarizing current injected into the amacrine cell were the same as the effects of sensitization. To capture the effects of either current, or sensitization, or both, we fit transformations consisting of horizontal shifting and scaling parameters, *T* = (*µ, ν*), between response curves in different conditions. To quantify the effects of sensitization, we computed the transformation, *T*_*Sens*_, that changed the control ganglion cell response curve from *G*_*L*_, during *L*_*late*_, to *G*_*E*_, during *L*_*early*_. To quantify the effects of hyperpolarizing current during *L*_*late*_, we computed the parameters, *T*_*Hyp*_, that transformed the ganglion cell response curve from *G*_*L*_ to *G*_*L−*_ during *L*_*late*_. We then tested whether a single transformation, *T*_*Both*_, fit to simultaneously capture the effects of hyperpolarizing current and sensitization could do as well as the two transformations, *T*_*Sens*_ or *T*_*Hyp*_, fit separately to capture sensitization and current injection, respectively. Because sensitization likely resulted from the effects of multiple amacrine cells, we computed a single parameter, *κ_−_*, to scale the effects of hyperpolarization (Figure 5A & B, see Experimental Procedures). Thus *T*_*Both*_ = (*µ, ν*) was fit to capture the effects of current from *G*_*L*_ to *G_L−_*, and simultaneously *κ*_−_ was fit so that *κ*_−_*T*_*Both*_ = (*κ_−_µ, κ_−_ν*) best captured the effects of sensitization from *G*_*L*_ to *G*_*E*_. We found that scaling the effects of hyperpolarizing current using the single transformation *κ_−_T*_*Both*_ captured 90.5 *±* 3.1% of the effect of sensitization as captured by the parameters *T*_*sens*_ fit only to sensitization (Figure 5C and S5A). Furthermore, the simultaneously fit transformation *T*_*Both*_ captured 98.4 *±* 7.2% of the effects of current as captured by *T*_*Hyp*_ fit only to current injection (Figure 5C and S5B).

**Figure 5:**
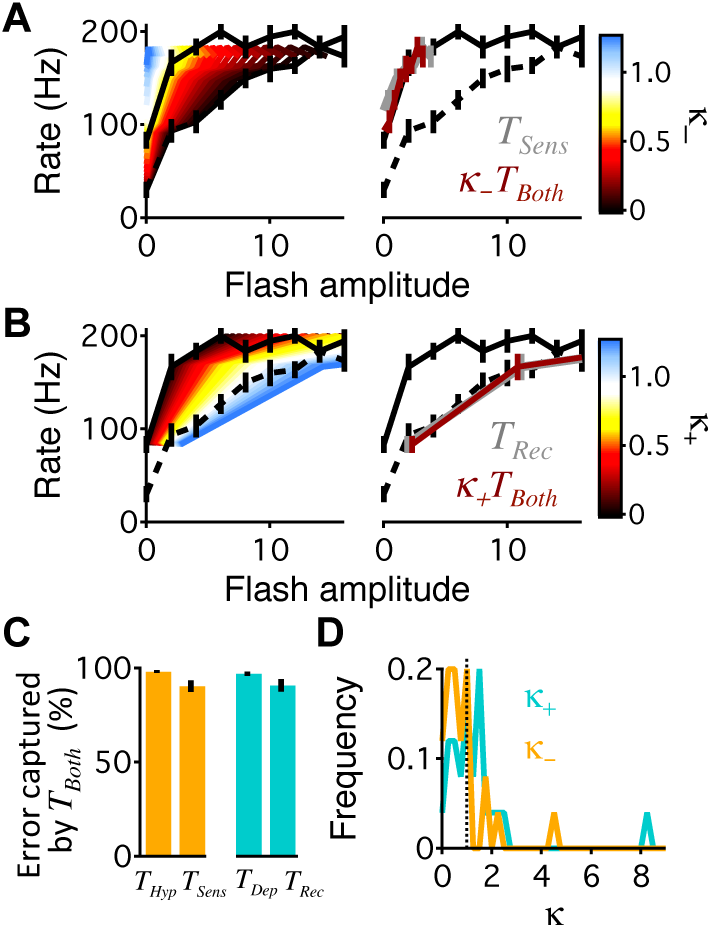
Amacrine cell transmission can create, or negate sensitization. (A) Left, ganglion cell response curves from an example cell under control conditions during *L*_*early*_, *G*_*E*_(*s*), (solid line) and during *L*_*late*_, *G*_*L*_(*s*) (dotted line). Curves in the background are scaled versions of the *κ*_−_*T*_*Both*_ = (*κ_−_µ_B−_, ν_B−_*) scaled by different values of *κ*. The curve with a value of *κ*_−_ = 1 indicates *T*_*Both*_, the best fit to transform *G*_*L*_(*s*) to the measured current injection curve, *G_L−_*. Right, Same measured control response curves, *G*_*E*_(*s*) and *G*_*L*_(*s*) along with the *L*_*late*_ curve, *G*_*L*_(*s*), transformed by two different transformations: *T*_*Sens*_ (grey, obscured by the maroon curve) and *κ*_−_*T*_*Both*_ (maroon), with optimized *κ_−_*. (B) Left, same as panel (A) for *κ*_+_*T*_*Both*_ = (*κ* + *µ_B_*_+_*, κ*_+_*ν_B_*_+_). Right, same as panel (A) for *T*_*Rec*_ (grey, obscured by the maroon curve) and *κ*_+_*T*_*Both*_(maroon), with optimized *κ*_+_. (C) Left, error captured by the simultaneous transformations *T*_*Both*_ and *κ*_−_*T*_Both_ as a percentage of the error captured by the separately fit transformations *T*_Hyp_ and *T*_*Sens*_, respectively. Right, the same for the simultaneous transformations *T*_*Both*_ and *κ*_+_*T*_*Both*_ and the single transformations *T*_*Dep*_ and *T*_*Rec*_. (D) Histograms of *κ* values for all pairs of cells that had ganglion cells that sensitized (adaptive index *>* 0.05) and were within 200 *µ*m from each other (n = 25 cell pairs, 19 amacrine cells) for the transformation of the hyperpolarizing curves during *L*_*late*_ (*κ*_*−*_) and depolarizing curves during *L*_*early*_ (*κ*_*−*_).

This indicated that amacrine hyperpolarization produced nearly the same type of effect on the ganglion cell light response as sensitization. The value of *κ*_*−*_ (Figure 5D) on average was 0.99 *±* 0.19, meaning that on average for the cell pairs we recorded from using 500*pA* pulses during *L*_*late*_ produced the same magnitude of effect as visual sensitization. We conclude that a decrease in inhibition from sustained amacrine cells is sufficient to cause sensitization of the ganglion cell light response.

We then computed whether depolarizing an amacrine cell and thus restoring the level of inhibition during *L*_*early*_ would cancel sensitization, as predicted by the model. We tested whether a single transformation, *T*_*Both*_, fit to simultaneously capture the effects of depolarizing current and the recovery from sensitization could do as well as the two transformations *T*_*Rec*_ or *T*_*Dep*_ fit separately to capture the recovery from sensitization and current injection, respectively. We found that the single transformation *κ*_+_*T*_*Both*_ = (*κ*_+_*µ, κ*_+_*ν*) captured 90.8 *±* 3.3% of the effect of the recovery from sensitization from *G*_*E*_ to *G*_*L*_ as captured by the parameters *T*_*Rec*_ fit only using *G*_*E*_ and *G*_*L*_ (Figure 5C and S5C). Furthermore, *T*_*Both*_ captured 97.1 *±* 1.1% of the effects of current as captured by *T*_*Dep*_ fit only to current injection using *G*_*E*_ and *G*_*E*+_ (Figure 5C and S5D). The value of *κ*_+_ (Figure 5D) was 1.49 ± 0.31, indicating that on average the effect of visual sensitization was 1.49 times larger than the effect of +500*pA* of current injected into a single amacrine cell during *L*_*early*_.

Experimentally decreasing inhibition from an amacrine cell during *L*_*late*_ produces a ganglion cell light response similar to the sensitized response during *L*_*early*_, and experimentally increasing inhibition from an amacrine during *L*_*early*_ cancels sensitization, producing a ganglion cell response similar to that during *L*_*late*_. We therefore conclude that decreased inhibition from sustained Off-type amacrine cells, which arises from the observed amacrine hyperpolarization (Figure 1G) and the observed decrease in amacrine transmission during sensitization (Figure 2C – E), is both necessary and sufficient to cause sensitization of the ganglion cell light response.

### 2.6 Direct amacrine cell current injection is sufficient to generate sensitization

Finally, we tested whether the signal within the amacrine cell itself is sufficient to sensitize nearby ganglion cells. We designed an experiment to present a low contrast visual stimulus while injecting current into an amacrine cell so only that cell experienced high contrast. The visual stimulus was comprised of exclusively low contrast (Figure 6A). To mimic the response to a high contrast visual stimulus, we used a record and playback approach (Manu and Baccus, 2011), periodically injecting a current that reproduced the voltage change in the amacrine cell recorded previously during high contrast (Figure 1E), while measuring the response of multiple ganglion cells. The high contrast current injected into the amacrine cell was sufficient to cause sensitization in ganglion cells (Figure 6B – D), and this effect was specific to nearby ganglion cells, having a space constant of *λ* = 150 ± 90*µ*m. Similar experiments in horizontal cells showed that the signal experienced by horizontal cells was not sufficient to cause sensitization (Figure 6D).

**Figure 6:**
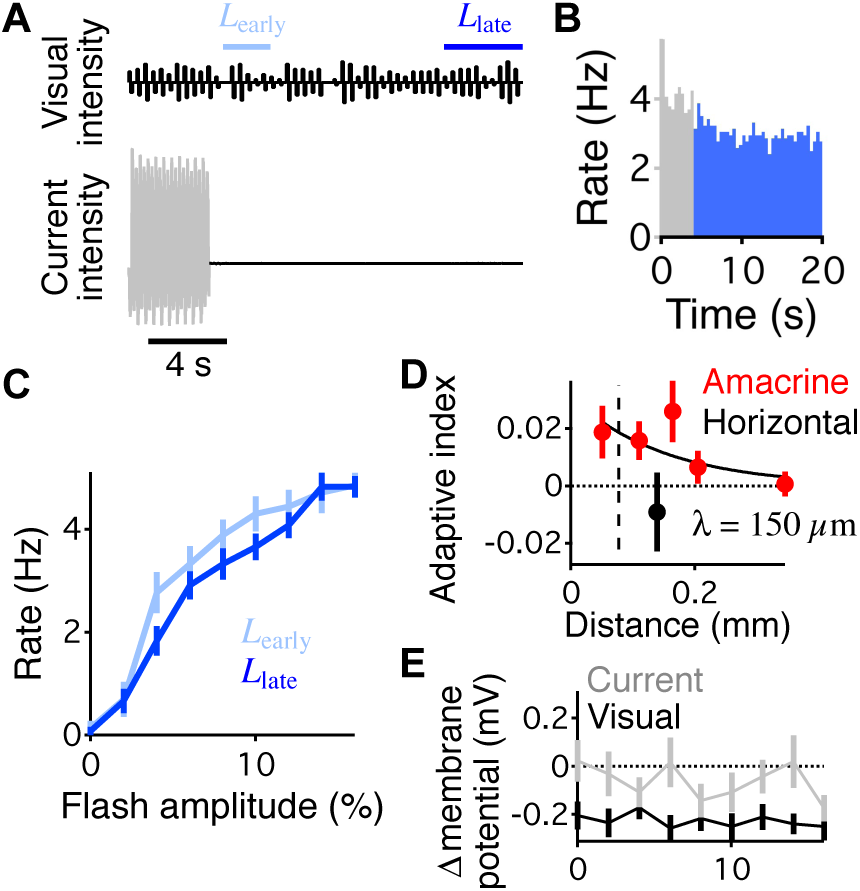
Current injection into amacrine cells is sufficient to sensitize ganglion cells. (A) The visual stimulus was a continuous low contrast composed of 9 different flashes randomly presented (top). Each flash lasted for 400 ms with 100 ms above the mean, 100 ms below the mean, and 200 ms at the mean intensity. Current was injected into a single amacrine cell to mimic its high contrast response using a record and playback approach (Manu and Baccus, 2011) (see Experimental Procedures). (B) Average response of a ganglion cell to the stimulus protocol in A. Grey indicates the time of current injection, and the blue color indicates the time without current injection. (C) Average response of the ganglion cell in B to the 9 different flashes presented in the visual stimulus during *L*_*early*_ and *L*_*late*_, as indicated in A. (D) Average adaptive index of all recorded fast Off ganglion cells plotted relative to the distance between the ganglion cells and amacrine (red) or horizontal cells (black). The adaptive index is 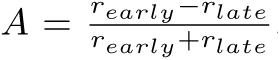, where *r*_early_ *r*_late_ are the average rate of the ganglion cell during *L*_*early*_ and *L*_*late*_, respectively. Data comes from a total of 76 amacrine and ganglion cell pairs and 10 horizontal and ganglion cell pairs. Black curve shows an exponential fit to the amacrine cell data. (E) Average difference in membrane potential between *L*_*late*_ and *L*_*early*_ for amacrine cells (n = 22) following a visual high contrast (black) or a high contrast current injected into the amacrine cell (grey). Negative values indicate that the cell was depolarized during *L*_*late*_ compared to *L*_*early*_. Error bars indicate s.e.m.

We examined the magnitude of sensitization caused by direct current injection into a single amacrine cell by calculating the shift in the ganglion cell response curve averaged across cell pairs. We found that visually induced sensitization was 9.1 ± 0.2 times greater for than for direct current injection into a single amacrine cell without high contrast visual stimulation. This difference likely occurs because the sensitizing field of ganglion cells is 300 – 410 *µ*m (Kastner and Baccus, 2013), considerably greater than the receptive field of a single amacrine cell (110 *µ*m), suggesting that multiple amacrine cells act during visual sensitization. Additionally, the effect of depression of amacrine cell transmission is expected to account for approximately 2/3 of the effect of sensitization, with the afterhyperpolarization of the amacrine cell membrane potential (Figure 1 & 4), accounting for the remainder. Although amacrine cells experienced an afterhyperpolarization during visual sensitization (Figure 1E), sensitization from direct current injection acts through an effect distinct from this afterhyperpolarization, since current injection did not consistently create an afterhyperpolarization in amacrine cells (Figure 6E). Thus, the signal experienced by single sustained Off amacrine cells during local high contrast is sufficient to sensitize nearby ganglion cells to a low contrast visual stimulus.

## 3 Discussion

Here we have shown that changes in inhibition in two sequential stages causes sensitization in retinal ganglion cells using neural recordings, dynamic perturbations, and computational modeling. A reduction of inhibition has been shown to underlie elevations of sensitivity in the visual cortex (Fu et al., 2014), and behavioral choice in drosophila (Jovanic et al., 2016) but how these computations unfold within the circuit is unknown. In taking advantage of known retinal pathways to dissect the computation of sensitization, we have focused in particular on changes in inhibitory transmission in the stage between amacrine cells and ganglion cells. Although the decrease in inhibition that underlies sensitization may derive from synaptic depression of the amacrine cell itself, it may also involve more complex polysynaptic transmission. Thus our studies reveal a mechanistic contribution from a particular component, the amacrine to ganglion cell stage, and further studies will be needed to further subdivide this stage into smaller mechanistic parts.

### 3.1 Adaptation arising from narrow-field amacrine cells

Our conclusion that sustained Off amacrine cells cause sensitization is consistent with a number of previous findings. Sensitization persists when the On pathway is blocked, making On type amacrine cells an unlikely candidate (Kastner and Baccus, 2013). A model that reproduces sensitization contains an inhibitory pathway with a tonic output that increases sensitivity during sensitization through disinhibition (Kastner and Baccus, 2011, 2013), and sustained Off amacrine cells have been shown to function largely through disinhibition (Manu and Baccus, 2011). Finally, sensitization occurs at a scale smaller than the ganglion cell receptive field center (Kastner and Baccus, 2011, 2013), making it less likely that the inhibitory cell will have a large receptive or projective field. Sustained Off amacrine cells are narrow field, having small receptive and projective fields (de Vries et al., 2011), especially when compared to transient cells like the polyaxonal amacrine cell (Cook et al., 1998; Baccus et al., 2008).

### 3.2 Establishing a mediator in a dynamic nonlinear system

Establishing that a mechanism is responsible for a function requires showing that the mechanism is necessary, sufficient, and is active at the correct time and place (Falkow, 2004; Kramer and Davenport, 2015). Although previous results show that transmission from GABAergic amacrine cells was necessary for sensitization, it was not known whether these cells mediated the process, or even whether stimulus evoked activity in these cells played a role. Our results show that a reduction in stimulus evoked amacrine cell inhibition is necessary, sufficient and correlated with changes in sensitization (Figure 7). The response to high contrast in sustained amacrine cells is correlated with (Figure 1 & 2) and sufficient to generate sensitization (Figure 6). Figure 5 shows that either the amacrine hyperpolarization or the reduction of transmission is necessary and sufficient to create sensitization. Taken together, these results show that sustained amacrine cells mediate retinal sensitization. In a nonlinear neural circuit whose state varies on a millisecond timescale, establishing causality requires both dynamic perturbations and measurements. In addition, it can be difficult to discern the right time and place that a mechanism could potentially cause the target effect, and several substantial difficulties face the rapidly growing field of neural perturbations to define neural mechanisms (Otchy et al., 2015; Phillips and Hasenstaub, 2016).

**Figure 7:**
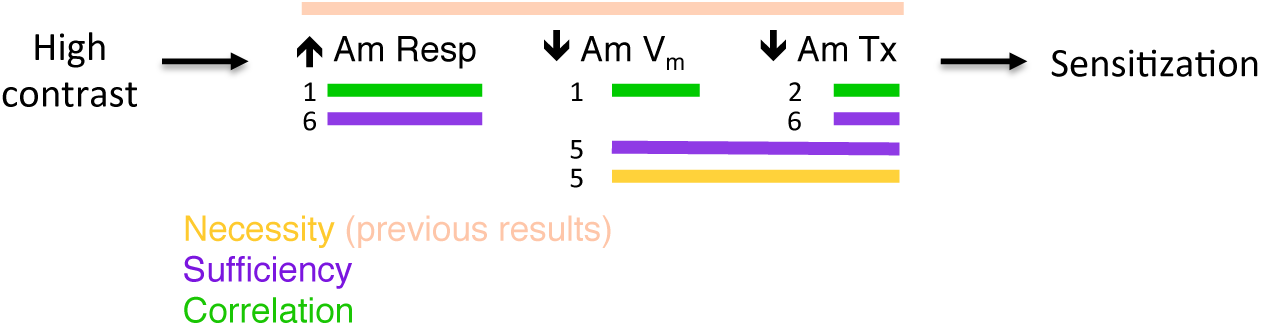
Summary of experimental findings. High contrast leads to an increase in amacrine response (Am Resp), a hyperpolarization of the amacrine cell membrane potential, and a decrease in amacrine transmission, which may be polysynaptic. Colored bars indicate different experimental conclusions. Previous results showed that GABAergic amacrine cells are necessary, but not whether they mediate the effect of sensitization. Numbers indicate figures that show the components are necessary, sufficient or correlated with sensitization. Overall, the amacrine cell hyperpolarization or transmission or both are necessary and sufficient for sensitization, and are correlated with sensitization.

One issue to consider is that the mechanism described is not just a particular neural component, such as a cell or synapse, but in fact the dynamic response of that component; some other dynamic response of the same component could cause a different process. Thus, tests of sufficiency and necessity must match the dynamic perturbation to the particular neural response pattern (Manu and Baccus, 2011), which requires both recording from and perturbing the same cells. Furthermore, showing that a mechanism is active at the right time and place must account for nonlinearities and delays in the circuit, and so a computational model of the system is critical in fulfilling this requirement, as it was in our study. Note that such a model need not account for all circuit details, but should account for the signals that flow through the perturbed neurons, and those flowing through other inaccessible neural pathways, as we have done here. Most current studies in the brain have not yet achieved these requirements, although sufficient experimental and computational methods exist, and one expects will be applied, to reveal the causal contribution of dynamic neural mechanisms of additional complex dynamic functions.

### 3.3 Updating of predictive sensitization

Sensitization has been proposed to form a prediction of the future location of a moving object, given the expectation that if an object moves, it will likely be present nearby in the near future (Kastner and Baccus, 2013). During visual sensitization, the effect of amacrine cell tranmission depression appears to have a larger effect on the ganglion cell response curve than does amacrine cell afterhyperpolarization (Figure 4). However, we found that directly manipulating the amacrine cell membrane potential from a single cell during sensitization could cancel most of the effect of sensitization (Figure 5). Although this difference could result from differences in experimental conditions in that in one case current was applied following a high contrast visual stimulus, it is worth considering that under natural conditions, a larger change in steady amacrine cell membrane potential might act to cancel sensitization due to a visual stimulus. If so, a new stimulus effective at depolarizing those amacrine cells could act at an immediate timescale—faster than sensitization—to potentially cancel previously created sensitization. This local cancellation of sensitization might occur even at the level of a single amacrine cell. Thus, given the predictive nature of sensitization to alert the ganglion cell to the potential for new motion of a localized object, a subsequent movement of that object that depolarizes Off-type amacrine cells could serve to cancel previous sensitization, thus both allowing for the response to new motion and acting to sensitize that new location for subsequent stimuli. Further experiments will be required to confirm this hypothesis of local stimuli to dynamically update the state of sensitization.

### 3.4 Adapting inhibition in two stages enables distinct sensitizing components

Adaptation opposes sensitization in the neural pathways that lead to a downstream neuron. Thus, the final result of net sensitization at any one time depends on there being an excess of sensitization over adaptation within those pathways. Under the model that decreasing excitation causes adaptation (Ozuysal and Baccus, 2012; Jarsky et al., 2011) and decreasing inhibition causes sensitization, net sensitization must arise from differences in the properties of excitatory and inhibitory plasticity. In the two components of sensitization we have identified—amacrine response adaptation and decreasing amacrine transmission— the particular dynamics, magnitudes and threshold of these processes will influence the amount of sensitization. Bipolar cells presumably cause amacrine response adaptation, and thus the properties of amacrine response adaptation and bipolar cell excitatory adaptation may directly cancel each other (Kastner and Baccus, 2013; Nagel et al., 2015), although differences may arise from different properties such as excitatory and inhibitory synaptic thresholds. In contrast, an advantage of generating sensitization in two stages may be that changes in amacrine transmission can allow sensitization to be generated with dynamics that are distinct from other excitatory plasticity in the circuit.

The properties of sensitization vary in their magnitude and time course between ganglion cells based upon the stimulus conditions (Kastner and Baccus, 2013). Generating a change in inhibition from the combined effects of two sources—putatively, bipolar synapses and amacrine synapses—could potentially allow for flexibility in the dynamics of sensitization in different ganglion cells. Furthermore, inhibition, even from sustained Off amacrine cells, plays multiple roles in the circuit (Manu and Baccus, 2011; de Vries et al., 2011), which likely places additional constraints upon how much any given element can adapt, and still maintain its broad functionality. Given that, as we have shown, inhibition influences the threshold of the ganglion cell response (Figures 2, 5 & 6), and that ganglion cell threshold is critical for the optimal response properties of the retina (Pitkow and Meister, 2012; Kastner et al., 2015), these amacrine cells might have further roles in the circuit that could place additional constraints on their response dynamics.

### 3.5 Changing inhibition as a form of short term information storage

Inhibitory plasticity has been proposed to support the excitatory-inhibitory balance seen in cortical circuits (Vogels et al., 2011; Landau et al., 2016). We have considered an alternative function, that sensitization arising from inhibitory plasticity acts as a form of short-term information storage over several seconds (Kastner and Baccus, 2013). The decrease in inhibition observed here accomplishes the same function as an increase in excitation as has been theorized in a computational model of cortical short-term working memory (Hempel et al., 2000; Mongillo et al., 2008; Pereira and Wang, 2015). If short-term synaptic plasticity does, in fact, underlie short-term memory, it should be considered that excitatory synaptic facilitation and inhibitory synaptic depression could both contribute, and that different dynamics could be combined from different types of synapses.

### 3.6 A general circuit for prediction

Sensitization is predictive in ganglion cells that act as feature detectors for stimuli such as differential motion. For a strong stimulus, detection of a cell’s preferred stimulus feature leads to enhanced sensitivity for that feature (Kastner and Baccus, 2013). Because adaptation and synaptic depression are highly prevalent in the nervous system, a general strategy for prediction may be to control the sensitivity for a feature by using an inhibitory pathway that suppresses that feature, allowing depression of inhibition to generate feature-specific sensitization, and thereby enhancing the future sensitivity for that feature. Enhanced sensitivity due to specific decrease in inhibition may therefore underlie sensitization observed in the cortex (Cohen-Kashi Malina et al., 2013; Ganmor et al., 2010; Mohar et al., 2013), and may reflect a general mechanism for prediction in the nervous system.

## 4 Experimental Procedures

### 4.1 Experimental preparation

Retinal ganglion cells of larval tiger salamanders of either sex were recorded using an array of 60 electrodes (Multichannel Systems) as described (Kastner and Baccus, 2011). A video monitor projected the visual stimuli at 30 Hz controlled by Matlab (Mathworks), using Psychophysics Toolbox (Brainard, 1997; Pelli, 1997). Stimuli had a constant mean intensity of 10*mW/m*^2^.

### 4.2 Intracellular recording

Simultaneous intracellular and multielectrode recordings from the isolated intact salamander retina were performed as described (Manu and Baccus, 2011). Sustained amacrine and horizontal cells were identified by their flash response and their spatiotemporal receptive fields (Figure 1C & S1A), with horizontal cells lacking an inhibitory surround and being greater than 300*µm* in diameter. For some of the cells we confirmed that they were in fact amacrine cells by filling and imaging them (n = 4) (Figure S1B).

The output of the ganglion cell response functions for the transmission experiments (Figure 2 & 5) was determined as the maximal firing rate in a 50 ms window averaged across all trials for each flash amplitude. All fast Off ganglion cells that responded in the control conditions (Figure 2B) with an average response over 11 Hz were included in the analysis to determine the changes in the transmission (Figure 2 & 5).

The record and playback experiments (Figure 6) were carried out as previously described (Manu and Baccus, 2011). Briefly, the response of an amacrine cell to a visual stimulus was recorded. Then the visual response during high contrast was deconvolved with an exponential filter, capturing the membrane time constant, to create the current necessary to inject to playback the response of the amacrine cell to high contrast. The injected current was scaled to have a standard deviation of 500 pA. This magnitude of current in sustained Off amacrine cells is estimated to cause a change in membrane potential with a standard deviation of *∼* 9.6*mV* (Manu and Baccus, 2011).

### 4.3 Receptive field measurement

Spatiotemporal receptive fields were measured in one or two dimensions by the standard method of reverse correlation (Chichilnisky, 2001) of the spiking response with a visual stimulus consisting of squares that were drawn from a binary distribution, or lines drawn from a Gaussian white noise distribution, such that

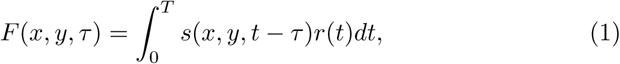

where *F* (*x, t, τ*) is the linear response filter at position (*x, y*) and delay *τ*, *s*(*x, y, t*) is the stimulus intensity at position (*x, y,*) and time *t*, normalized to zero mean, *r*(*t*) is the firing rate of a ganglion cell or the membrane potential of an amacrine cell, and *T* is the duration of the recording.

For all amacrine cell recordings, the receptive field of the amacrine cell was measured online, and the stimulus was centered onto the receptive field of the amacrine cell. The distance between an amacrine and ganglion cell was determined as the distance between the centers of two-dimensional Gaussian fits to their receptive fields.

### 4.4 Measuring changing response curves

During high contrast the amplitude of the flash flickered with a period of 200 ms and 100% Michelson contrast, defined as (*I_max_ − I_min_*)*/*(*I*_*max*_ + *I*_*min*_). During low contrast the contrast amplitude of the flash varied randomly every 400 ms, and was one of nine values that ranged from 0 – 16% contrast. Changing the distribution of amplitudes slower than the integration time of ganglion cells allowed for a rapid measurement of the ganglion cell response function without having to also measure the ganglion cell temporal filter (Brenner et al., 2000). Synchronized to the visual stimulus, we injected, randomly interleaved, hyperpolarizing or depolarizing current pulses into the amacrine cell. We computed response curves that were a function of the visual stimulus during *L*_*early*_, *G*_*E*_(*s*), as a control, *G*_*E*+_(*s*) using depolarizing current, and *G*_*E−*_(*s*) using hyperpolarizing current, and similarly, *G*_*L*_(*s*), *G*_*L*+_(*s*), and *G*_*L−*_(*s*) during *L*_*late*_ (Figure 2A & B).

We examined the effect of amacrine cell current on the control ganglion cell light response during *L*_*late*_, *G*_*L*_(*s*), which approximated the steady state low contrast response. The response curves under different stimulus conditions were approximately sigmoidal but with different slopes and midpoints, suggesting that one curve could be transformed into another by four parameters that shifted and scaled the response curve horizontally or vertically, or both. Therefore, we captured the transformation caused by amacrine current with manipulations to *G*_*L*_(*s*) reflecting changes in four parameters. The two parameters that horizontally shifted and scaled were *µ*, which shifted *G*_*L*_(*s*) with respect to the visual stimulus *s*, and *ν*, which scaled *G*_*L*_(*s*) horizontally with respect to *s*. These two parameters changed the threshold and slope of *G*_*L*_(*s*). The two parameters that vertically shifted and scaled the curve were *α*, which shifted the output vertically, and *β*, which scaled the output. These two parameters change the minimal and maximal output of the response function (Figure S2A).

Previously it was concluded that current injection into sustained Off amacrine cells changed the fast Off ganglion cell light response consistent with a presynaptic effect delivered prior to the sharp threshold at the bipolar cell terminal (Manu and Baccus, 2011). Therefore, we measured how much of the effect of the current could be captured by only using the input parameters, *µ* and *ν* for the transformation, compared to using all four parameters. As an example using the depolarizing current condition: for the transformation using all four parameters, *T*_4*Dep*_ = (*µ*_+*I*_, ν_+*I*_, α_+*I*_, β_+*I*_), this was computed by finding the parameters that minimized the difference between *G*_*L*_(*s*) and *βG*_*L*+_(*e^ν^ s* + *µ*) + *α*. Whereas, for the transformation that only manipulated the input parameters, *T*_*Dep*_ = (*µ*_+*I*1_, ν _+*I*_), we sought to find the values that minimized the difference between *G*_*L*_(*s*) and *G*_*L*+_(*e^ν^ s* + *µ*). The multiplicative scaling parameter *ν* was raised to an exponent so that cells would exert their multiplicative effects independently, i.e. one amacrine cell acting with a scaling parameter of *ν* would scale the stimulus by *e*^*ν*^, and two cells would scale the stimulus by *e*^2ν^.

For hyperpolarizing current during *L*_*early*_, which changed the response curve from *G*_*L*_(*s*) to *G_L−_*(*s*), the two parameters of the transformation *T*_*Hyp*_ = (*µ_−I_, ν*_*−I*_) accounted for 79.9 *±* 3.2% of the total effect of current injection, determined by using the transformation *T*_4*Hyp*_. In solving for *T*_*Hyp*_, we constrained *µ* and *ν* to be positive. For depolarizing current, which changed the response curve from *G*_*L*_(*s*) to *G*_*L*+_(*s*), the transformation *T*_*Dep*_ accounted for 82.3*±*3.2% of the total effect of current injection, *T*_4*Dep*_. For the transformation *T*_*Dep*_, we constrained *µ* and *ν* to be negative. Therefore, a transformation of the input dependence of the light response captures most of the effect of current injection, consistent with current injection predominantly working through an effect on the bipolar cell terminal, presynaptic to fast Off-type ganglion cells (n = 60 cell pairs, 24 amacrine cells).

We also measured how much of sensitization could be captured by only the two parameters of the transformation *T*_*Sens*_ = (*µ_S_, ν_S_*). For sensitization, which changes the light response from *G*_*L*_(*s*) during *L*_*late*_ to *G*_*E*_(*s*) during *L*_*early*_, a horizontal shifting and scaling by *T*_*Sens*_ accounted for 83.2 *±* 2.8% of the total effect of sensitization, found using *T*_4Sens_ = (*µ_S_, ν_S_, α_S_, β*_*S*_). For the recovery from sensitization, changing *G*_*E*_(*s*) to *G*_*L*_(*s*) using only a horizontal shifting and scaling, *T*_*Sens*_, accounted for 87.8 ± 2.0% of the total effect of recovery from sensitization. Thus, most of the effect of sensitization was also captured by a horizontal shifting and scaling of the light response, consistent with a presynaptic effect (n = 43 cell pairs, 23 amacrine cells).

All ganglion response curves were fit using a sigmoid function:

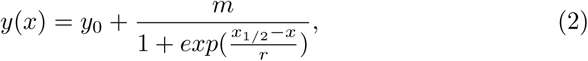

where *m* + *y*_0_ is the maximum value, *x*_1*/*2_ is the x value at which the function equals half the distance between the maximum value and *y*_0_, and *r* controls the rate at which the function increases. The curves were not extrapolated beyond the data limits. The error metric was the root mean squared (rms) difference between the sigmoid fits of the measured response function and the transformed response function. The fraction of the response that was captured by a two-parameter transformation *T*_2_ versus the *T*_4_ transformation was computed as 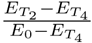; where *E*^*T2*^ and *E*^*T4*^ were the errors remaining after the *T*_2_ and *T*_4_ transformations, respectively, and *E*_0_ is the rms difference between the curves without any manipulation.

### 4.5 Scaling parameter *κ* to account for multiple amacrine cells and the magnitude of the current injection

To account for the effects of multiple amacrine cells and the fact that the effect of current injection on a single cell may differ in magnitude from the effect of sensitization, we computed for the effect of current injection the parameter *µ* that horizontally shifted the ganglion cell response curve, and the parameter *ν* that horizontally scaled the ganglion cell response curve, *G*(*e^ν^ s*+*µ*) to transform the ganglion cell response curve during the control condition onto the response curve under current injection, yielding the transformation *T*_*Hyp*_ = (*µ_−I_, ν_−I_*) for the transformation of the control curve, *G*_*L*_(*s*), onto the hyperpolarized curve, *G_L−_*(*s*), during *L*_*late*_. For comparison, we computed the effects of sensitization between *G*_*L*_(*s*) during *L*_*late*_ and *G*_*E*_(*s*) during *L*_*early*_ as the transformation *T*_*Sens*_ = (*µ_S_, ν_S_*). We then fit all parameters, (*µ_B−_, ν_B−_, κ*), together so that the transformation *T*_*Both*_ = (*µ_B−_, ν_B−_*) would most closely match the effects of current by transforming *G*_*L*_(*s*) to approximate *G_L−_*(*s*) and so that the scaled transformation *κ*_−_*T*_*Both*_ = (*κ_−_µ_B−_, κ_−_ν_B−_*) would most closely match the effects of sensitization by transforming *G*_*L*_(*S*) to approximate *G*_*E*_(*s*). The same was done for the comparison of *κ*_+_*T*_*Both*_, *T*_*Rec*_ and *T*_*Dep*_. The fraction of error captured by the single transformation *T*_*Both*_ compared to the transformation *T*_*Sens*_ was measured as 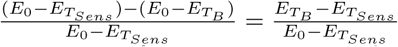; where *E*_*T*__*B*_ is the error for *T*_*Both*_, *E*_*TSens*_ is the errors for *T*_*Sens*_, and *E*_0_ is the rms difference between the *G*_*L*_(*s*) and *G*_*E*_(*s*) curves without any manipulation. The fraction of error captured by *T*_*Both*_ compared to the transformations *T*_*Hyp*_, *T*_*Rec*_ and *T*_*Dep*_ was similarly computed.

### 4.6 Linear-Nonlinear-Kinetic model

The LNK model consisted of two pathways, each consisting of a linear temporal filter *F*_*LNK*_(*t*), a static nonlinearity, *N* (*g*), and a first order kinetic system defined by a transition matrix **Q**(*u*). This model was optimized as described (Ozuysal and Baccus, 2012), and summarized here, with the exception that the output of the first inhibitory pathway was joined with the second excitatory pathway prior to the threshold. The components were parameterized as described below, and all parameters were fit together using a constrained optimization algorithm. For each pathway, the stimulus, *s*(*t*), was passed through a linear temporal filter, *F*_*LNK*_(*t*), and a static nonlinearity, *N* (*g*),

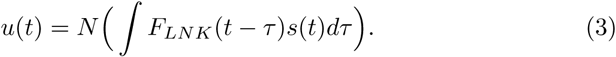

Although these two initial stages have the same structure as the linear-nonlinear (LN) model, the filter and nonlinearity are different functions than those computed for an LN model, and are optimized, rather than computed using reverse correlation. The kinetics block of the model was a Markov process defined by

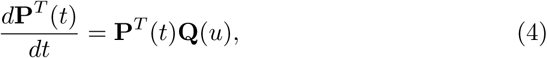

where **P**(*t*) is a column vector of *m* fractional state occupancies such that Σ_*i*_ *P*_*i*_ = 1, and **Q** is an *m × m* transition matrix containing the rate constants *Q*_*ij*_ that control the transitions between states *i* and *j*, with *Q*_*ii*_−Σ_*i*≠*j*_ *Q*_*ij*_. After this differential equation was solved numerically, the output of the model, *r*′(*t*) was equal to one of the state occupancies scaled to a response in millivolts,

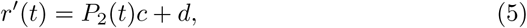

where *c* and *d* are a scaling and offset term for the entire recording. States and rate constants were defined as,

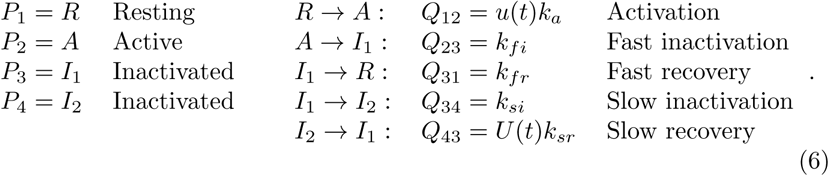

The change in state occupancy was thus determined as

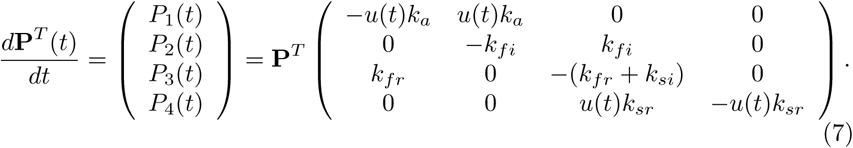

A more complex model with an additional adapting stage in the inhibitory pathway representing bipolar cell inputs to the amacrine cell would be needed capture the two-stage adaptation in the inhibitory pathway seen for a localized stimulus. This model for a spatiotemporal stimulus would require many additional parameters for spatiotemporal filters for a population of amacrine cells and an additional adaptive stage, and thus would not be well constrained with current measurements.

### 4.7 Contribution of amacrine cell membrane potential adaptation to sensitization

To determine the contribution of the observed amacrine cell membrane potential adaptation to sensitization (Figure 4), we first found the equivalent offset for the LNK model that matched the measured amacrine cell membrane potential difference between *L*_*early*_ and *L*_*late*_ (Figure 1G). We stimulated the model with the identical stimulus presented to the amacrine cells (Figure 1E), and then normalized the standard deviations of the data to be the same as in the model (Figure S4). This yielded the normalized difference in baseline membrane potential between *L*_*early*_ and *L*_*late*_ recorded from amacrine cells (Figure 1G), which was used as the model offset equivalent to the measured hyperpolarization.

To then determine the effect of amacrine membrane potential adaptation to sensitization, we computed an LN model fit to the LNK model that contained a single nonlinear response curve shown in Figure 4. To compute the LN model, we stimulated the model with high contrast (50%) flashes followed by low contrast Gaussian white noise, either with or without the normalized membrane potential offset applied to the inhibitory pathway. The low contrast white noise was chosen to create a response in the models inhibitory pathway that most closely matched the inhibitory response in the model during the low contrast flash protocol of Figure 1E.

### 4.8 Comparison of sensitization from visual stimulation and current

We performed a bootstrap analysis, where we sampled with replacement from the 57 ganglion cells that sensitized to the high contrast visual stimulations (i.e. adaptive index *>* 0) and for which we also had responses in those ganglion cells to the high contrast current injection into an amacrine cell. For each resampling we found the shifting along the x-axis that minimized the difference between the *L*_*early*_ and *L*_*late*_ curves, and took the average ratio of that scaling between visual high contrast stimulation and current injection.

## Acknowledgements

We thank D. Baylor, P. Jadzinsky, F. Zenke, S. Mensi, C. Pozzorini, and F. Dunn for helpful discussions, and J. Raymond for commenting on the manuscript. This work was supported by grants from the NEI, Pew Charitable Trusts, McKnight Endowment Fund for Neuroscience, the Alfred P. Sloan Foundation and the E. Matilda Ziegler Foundation (S.A.B.); by the Stanford Medical Scientist Training Program, an NSF IGERT graduate fellowship, and an NIH R25 (R25MH060482) (D.B.K).

## Author Contributions

D.B.K. and S.A.B. designed the study, D.B.K performed the experiments, D.B.K. and S.A.B. developed the analyses, D.B.K. and G.P. processed the cells for imaging, Y.O. fit the LNK model, and D.B.K and S.A.B. wrote the manuscript.

---

**Figure S1:**
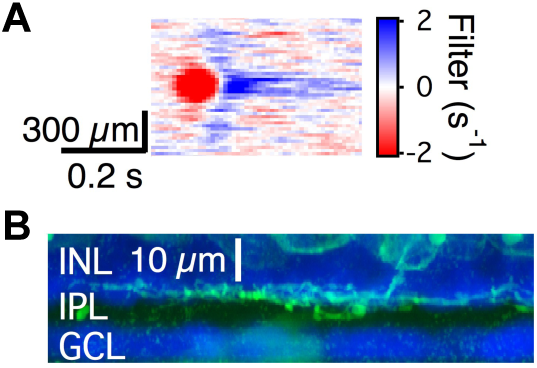
Amacrine cell identification. (A) Spatiotemporal receptive field of an amacrine cell. Stimulus intensities are indicated by the color scale, with red indicating an intensity below the mean value. (B) Image of the amacrine cell from A. INL is the inner nuclear layer, IPL is the inner plexiform layer, and GCL is the ganglion cell layer.

**Figure S2:**
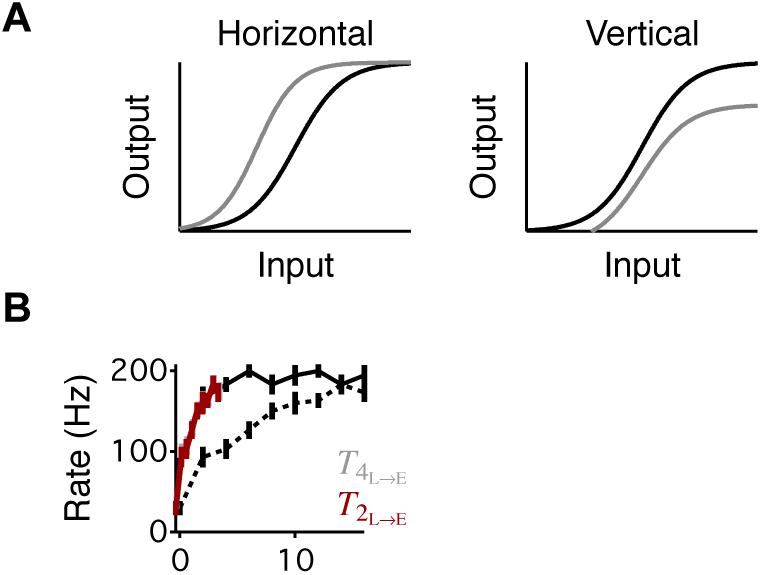
Amacrine current effects on the ganglion cell response function. (A) Example transformation using only a horizontal axis shifting and scaling (left), and using only the vertical axis parameters (right). (B) Same cell as in Figure 2B, with two transformations of *L*_*late*_ onto *L*_*early*_. The first transformation, *T*_4_ uses all 4 parameters (grey, obscured by maroon line), while the second transformation *T*_2_ only uses the horizontal axis parameters (see Experimental Procedures).

**Figure S3:**
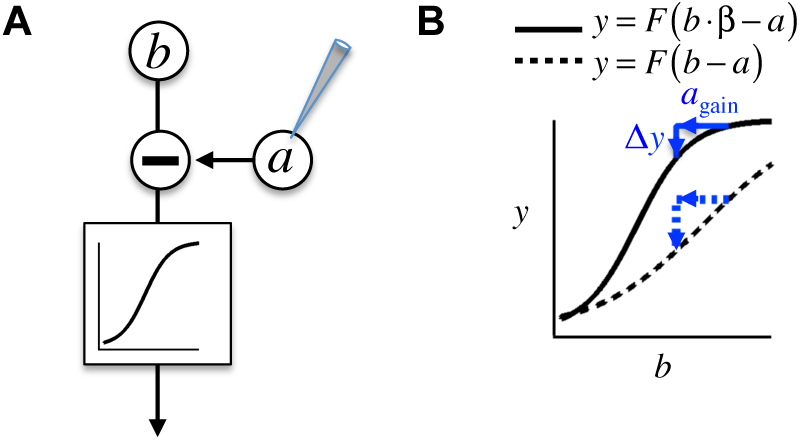
Example interaction between neurons in a nonlinear pathway. (A) Schematic diagram of an excitatory cell *b* and an inhibitory cell *a* that acts prior to a nonlinear function *F* () and is under direct experimental manipulation. (B) Diagram of the output of the system in two conditions, when the gain *β* of *b* is high (solid line) and when the gain is low (dotted line). In this example the output neuron *a* causes an effect ∆*y* when it is perturbed. In the two conditions, even though the gain of *a* is unchanged, it has a different effect because the resting condition is at a different point of the response curve.

**Figure S4:**
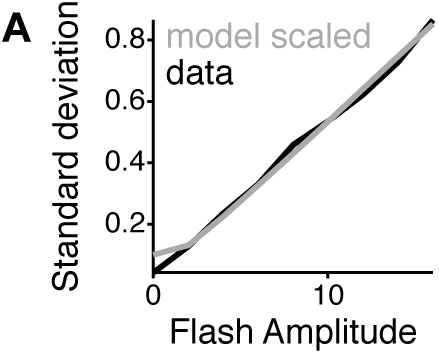
Scaling of LNK model to match data. Scaled standard deviation of the inhibitory pathway of the LNK model (Figure 3B) at the point following the inhibitory filter as a function of flash amplitude. This value is compared to the average standard deviation of the amacrine cell responses to the same stimulus from the amacrine cells in Figure 1E.

**Figure S5:**
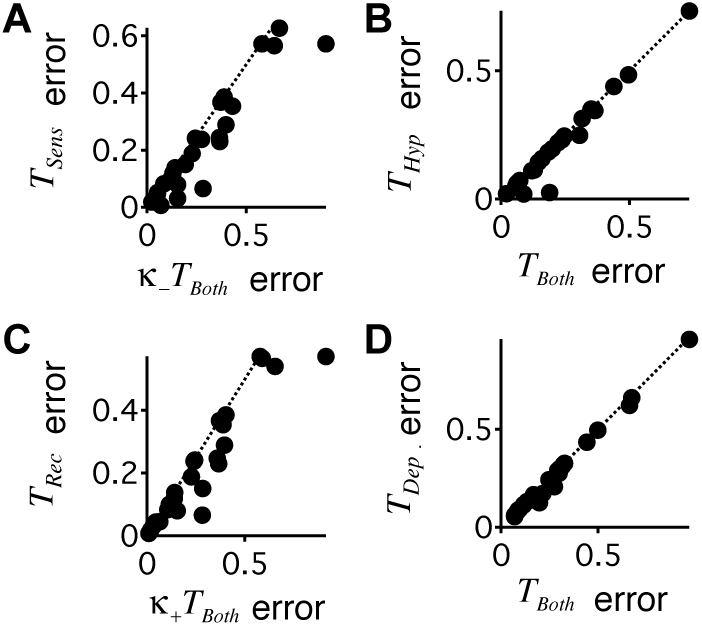
Comparison of error for transformations from current injection and the scaled effects of sensitization. (A) Error using the transformation *T*_*Sens*_ (ordinate) fit only to capture the changes from *G*_*L*_(*s*) to *G*_*E*_ (*s*) produced by sensitization, compared to the error from transformation *κ_−_T_Both_*(*κ_−_µ_B−_, κ_−_ν_B−_*) (abscissa) fit simultaneously to capture the changes from sensitization (*G*_*L*_ (*s*) to *G*_*E*_ (*s*)) and the changes from hyperpolarizing current injection (*G*_*L*_ (*s*) to *G_L−_*). The error represents the fraction of the change in rms response unaccounted for by the transformation. Each point is a cell pair. (B) Error using the transformation *T*_Hyp_ (ordinate) fit only to capture the changes from *G*_*L*_(*s*) to *G*_*L−*_ produced by hyperpolarizing current injection during *L*_*late*_, compared to the error from transformation *T*_*Both*_ = (*µ_B−_, ν_B−_*) (abscissa) fit simultaneously to capture both sensitization (*G*_*L*_(*s*) to *G*_*E*_ (*s*)) and current injection (*G*_*L*_ (*s*) to *G*_*L*−_). (C) Same as panel (A) for depolarizing current injection, and the transformations *T*_*Rec*_ (ordinate), fit only to capture the changes the recovery from sensitization (*G*_*E*_(*s*) to *G*_*L*_(*s*)), and the simultaneously fit *κ*_+_*T*_*Both*_(*κ*_+_*µ_B_*_+_*, κ*_+_*ν*_*B*+_) transformation to capture both the effects of depolarization and the recovery from sensitization. (D) Same as panel (B) for the transformations *T*_*Sens*_, fit only to capture the effects of depolarizing current during *L*_*early*_ (ordinate) and *T*_*Both*_ = (*µ*_*B*+_*, ν*_*B*+_), fit simultaneously to capture the effects of depolarizing current and the recovery from sensitization.

